# Spinal astrocyte-neuron lactate shuttle contributes to the PACAP/PAC1 receptor-induced nociceptive behaviors

**DOI:** 10.1101/317156

**Authors:** Yuki Kambe, Masafumi Yokai, Ichiro Takasaki, Takashi Kurihara, Atsuro Miyata

## Abstract

Previously, we showed that spinal pituitary adenylate cyclase-activating polypeptide (PACAP)/PAC1 receptor signaling triggers long-lasting pain behaviors through astroglial activation. Since astrocyte-neuron lactate shuttle (ANLS) could be essential for long-term synaptic facilitation, we aimed to elucidate a possible involvement of spinal ANLS in the development of the PACAP/PAC1 receptor-induced pain behaviors. A single intrathecal administration of PACAP induced short-term spontaneous aversive behaviors, followed by long-lasting mechanical allodynia. These pain behaviors were inhibited by DAB, an inhibitor of glycogenolysis, and this inhibition was reversed by simultaneous L-lactate application. In the cultured spinal astrocytes, the PACAP-evoked glycogenolysis and lactate secretion were inhibited by a protein kinase C (PKC) inhibitor, and the PKC inhibitor attenuated the PACAP-induced pain behaviors. Finally, an inhibitor for the monocarboxylate transporters blocked the lactate secretion from the spinal astrocytes and inhibited the PACAP-induced pain behaviors. These results suggested that PAC1 receptor-PKC-ANLS signaling is involved in the PACAP-induced pain behaviors.

## Introduction

Transfer of lactate from astrocytes to neurons is activated when synaptic activity is increased, and this mechanism is now known as the astrocyte-neuron lactate shuttle (ANLS), that could account for the coupling between synaptic activity and energy delivery (***Magistretti and Allaman, 2015***). Since lactate secretion from astrocytes is mainly originated from glycogen via glycogenolysis rather than from glycolysis (***Dringen et al., 1993***), inhibition of glycogenolysis or lactate transport in rat hippocampus significantly attenuated the working memory and long-term fear memory (***Newman et al., 2011; Suzuki et al., 2011***). In addition, a brain-specific glycogen synthase knockout mouse showed a significant deficiency in the acquisition with an associative learning task and concomitantly, activity-dependent changes in hippocampal synaptic strength (***Duran et al., 2013***). It is also reported that ANLS in amygdala is critical for the reconsolidation of cocaine memory in rats (***Boury-Jamot et al., 2015; Zhang et al., 2015***). At the cellular level, lactate may potentiate NMDA receptor-mediated currents in cultured neurons and neurons in the nucleus of the solitary tract (***Nagase et al., 2014; Yang et al., 2014***). These findings suggested that ANLS could contribute to many forms of neuronal plasticity in the CNS, including plastic changes in the spinal nociceptive transmission. However, it is still unclear what kinds of neurotransmitters evoke the ANLS activation during any forms of neuronal plasticity.

Pituitary adenylate cyclase-activating polypeptide (PACAP) was originally isolated from ovine hypothalamic extracts based on its ability to stimulate adenylate cyclase in rat anterior pituitary cell cultures (***Miyata et al., 1989; Miyata et al., 1990***). In normal state, PACAP specific receptor, PAC1 receptor, is particularly abundant in central nervous system (CNS) including spinal dorsal horn (***Dickinson et al., 1999;Jongsma et al., 2000;Sakashita et al., 2001;Vaudry et al., 2009; Yokai et al., 2016***), where PACAP-immunoreactive fibers are also considerably localized (***Moller et al., 1993; Dun et al., 1996a; Dun et al., 1996b; Narita et al., 1996***), and PACAP mRNA/immunoreactivity in rat dorsal root ganglia is markedly upregulated in peripheral nerve injury or inflammation (***Zhang et al., 1995; Zhang et al., 1998; Jongsma et al., 2003; Mabuchi et al., 2004***). These observations coupled with other lines of evidence propose that PACAP/PAC1 receptor system could play an important role in the modulation of spinal nociceptive transmission.

We have previously demonstrated in mice that a single intrathecal (i.t.) injection of PACAP or a PAC1 receptor specific agonist, maxadilan (Max) (***Moro and Lerner, 1997***), induced spontaneous aversive behaviors, such as licking, biting, and scratching directed toward the caudal part of the body for more than 30 min (***Shimizu et al., 2004; Ohnou et al., 2016***), and the aversive behaviors were followed by a induction of the mechanical allodynia, which lasted for more than 84 days (***Yokai et al., 2016***). However, vasoactive intestinal polypeptide (VIP), which share VIP/PACAP (VPAC) 1 or VPAC2 receptors with PACAP, failed to induce these nociceptive behaviors (***Ohnou et al., 2016; Yokai et al., 2016***). Co-treatment of a PAC1 receptor antagonist, max.d.4 with PACAP, almost completely inhibited the induction of both aversive behaviors and the mechanical allodynia (***Ohnou et al., 2016; Yokai et al., 2016***). These results suggested critical role of PAC1 receptor in nociceptive transmission. Immunohistochemical and immunoblotting studies revealed that spinal application of PACAP or Max induced upregulation of an astrocyte marker, glial fibrillary acidic protein (GFAP) at least for 84 days, and L-α-aminoadipate, an astroglial toxin, attenuated the induction of both aversive behaviors and mechanical allodynia. Interestingly, L-α-aminoadipate were still effective to reverse the mechanical allodynia even if it was i.t.- administered 84 days after spinal PAC1 receptor stimulation (***Yokai et al., 2016***). These results suggest that long-lasting spinal astocytic activation underlies the PACAP/PAC1 receptor-induced aversive behaviors and mechanical allodynia. However, how spinal astrocytes contribute to these nociceptive behaviors is still unknown.

Glycogen is the single largest energy reserve in the brain, and is predominantly localized in astrocytes (***Cataldo and Broadwell, 1986***). Among the neurotransmitters and bioactive substances examined up until today, PACAP is suggested to have potent glycogenolytic activity with an EC50 value of 0.08 nM (***Magistretti et al., 1998***). To the best of our knowledge, PACAP has the highest efficacy to induce glycogenolysis. However, it is not known whether PACAP is involved in the spinal ANLS. Therefore, we aimed to examine possible involvement of PACAP/PAC1 receptor system in the spinal ANLS, and also test whether the spinal ANLS contributes to the PACAP/PAC1 receptor-induced nociceptive behaviors.

## Results

### PACAP/PAC1 receptor-induced nociceptive behaviors were attenuated by the inhibition of glycogen phosphorylase with DAB

We first examined whether spinal ANLS played important role in the PACAP/PAC1 receptor-induced nociceptive behaviors in mice. After a single i.t. administration of Max (50 pmol) or PACAP (100 pmol), aversive behaviors such as licking and biting gradually appeared within 0 ~ 5 min, reached to the plateau around 15 ~ 20 min, and maintained at least for 30 min. The co-injection of DAB, a glycogen phosphorylase (PYGB) inhibitor, dose-dependently (1 ~ 100 pmol) attenuated the development of the Max- or PACAP-induced aversive behaviors (***Figure 1A, C***). In addition, a single i.t. administration of Max or PACAP markedly decreased mechanical threshold (induction of mechanical allodynia) from day 1 (after the cessation of the aversive behaviors), and this decrease persisted at least for 21 days after the administration. The co-injection of DAB with Max or PACAP also dose-dependently attenuated the induction of mechanical allodynia by Max or PACAP (***Figure 1B, D***).

**Figure 1.**
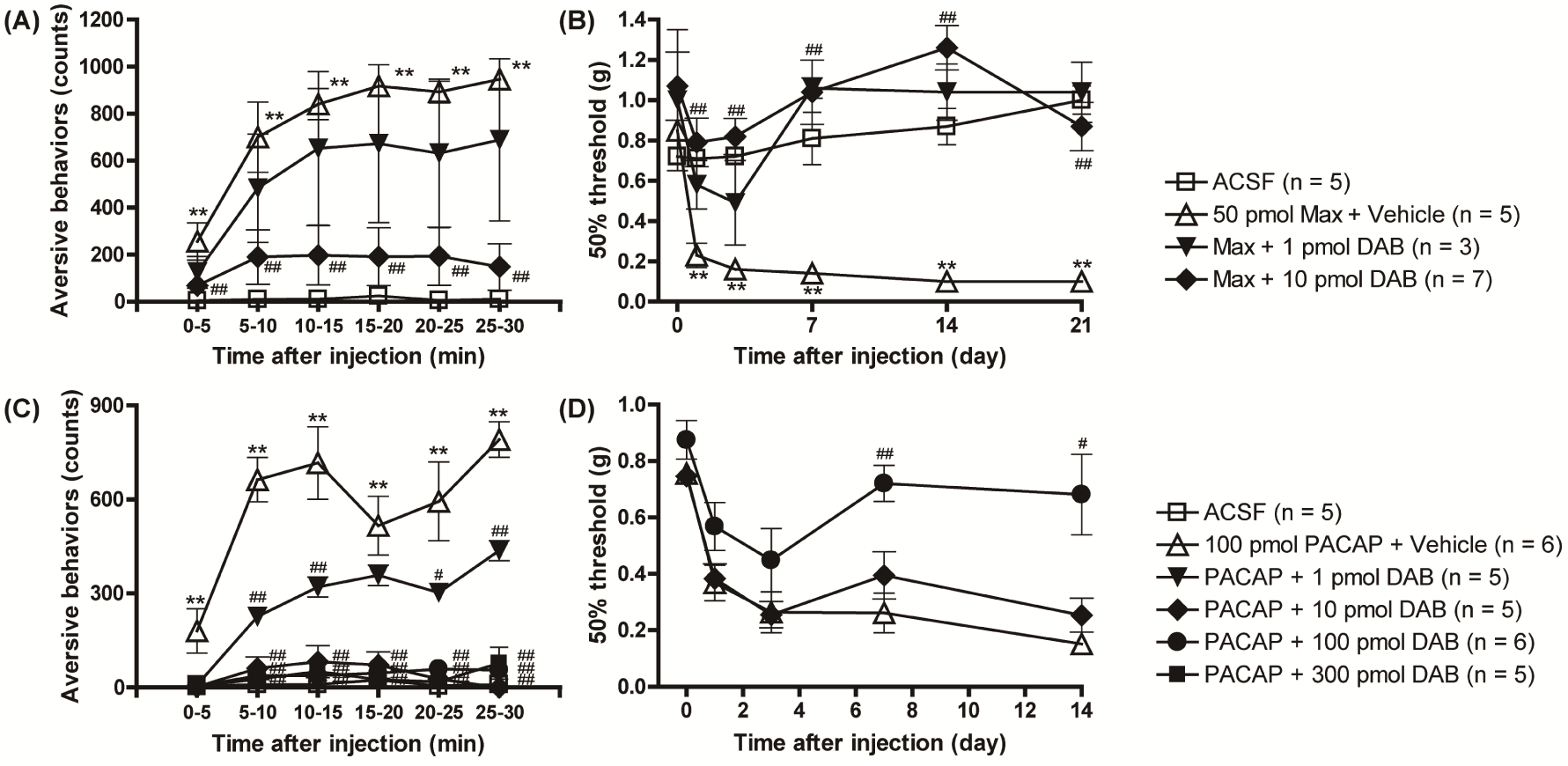
A critical role of glycogenolysis in PACAP/PAC1 receptor-evoked nociceptive behaviors. Simultaneous i.t. injection of DAB (1 or 10 pmol), a glycogenolysis inhibitor, with Max (50 pmol), blocked the induction of aversive behaviors (**A**) and mechanical allodynia (**B**). Simultaneous i.t. injection of DAB (1 ~ 300 pmol) with PACAP (100 pmol) prevented the induction of aversive behaviors (**C**) and mechanical allodynia (**D**). *P<0.05 and **P<0.01 when compared with ACSF data. ^#^P<0.05 and ^##^P<0.01 when compared with Max + Vehicle in (A) and (B) or PACAP + Vehicle in (C) and (D). Statistical significance was evaluated by the Tukey test for (A) and (C), and Steel test for (B) and (D).

### Suppression of the PAC1 receptor-evoked nociceptive behaviors by DAB was reversed by i.t. co-injection of L-lactate

Because lactate secretion from astrocytes was mainly derived from glycogen store (***Dringen et al., 1993***), we then asked whether the inhibition of the PAC1 receptor-induced nociceptive behaviors by DAB could be reversed by the co-administration of exogenous L-lactate. I.t. injection of L-lactate (1 nmol) in combination with Max (50 pmol) + DAB (10 pmol) significantly reversed the anti-aversive and anti-allodynic effects of DAB on the PAC1 receptor-induced nociceptive behaviors (***Figure 2A, B***). Similarly, i.t. supplementation of L-lactate (1 nmol) also blocked the inhibitory effects of DAB (100 pmol) on the PACAP-induced aversive behaviors and mechanical allodynia (***Figure 2C, D***). These results suggested that spinal ANLS has critical role in the PACAP/PAC1 receptor-induced nociceptive behaviors. However, L-lactate (10 nmol) alone did not show any pain related behaviors (data not shown).

**Figure 2.**
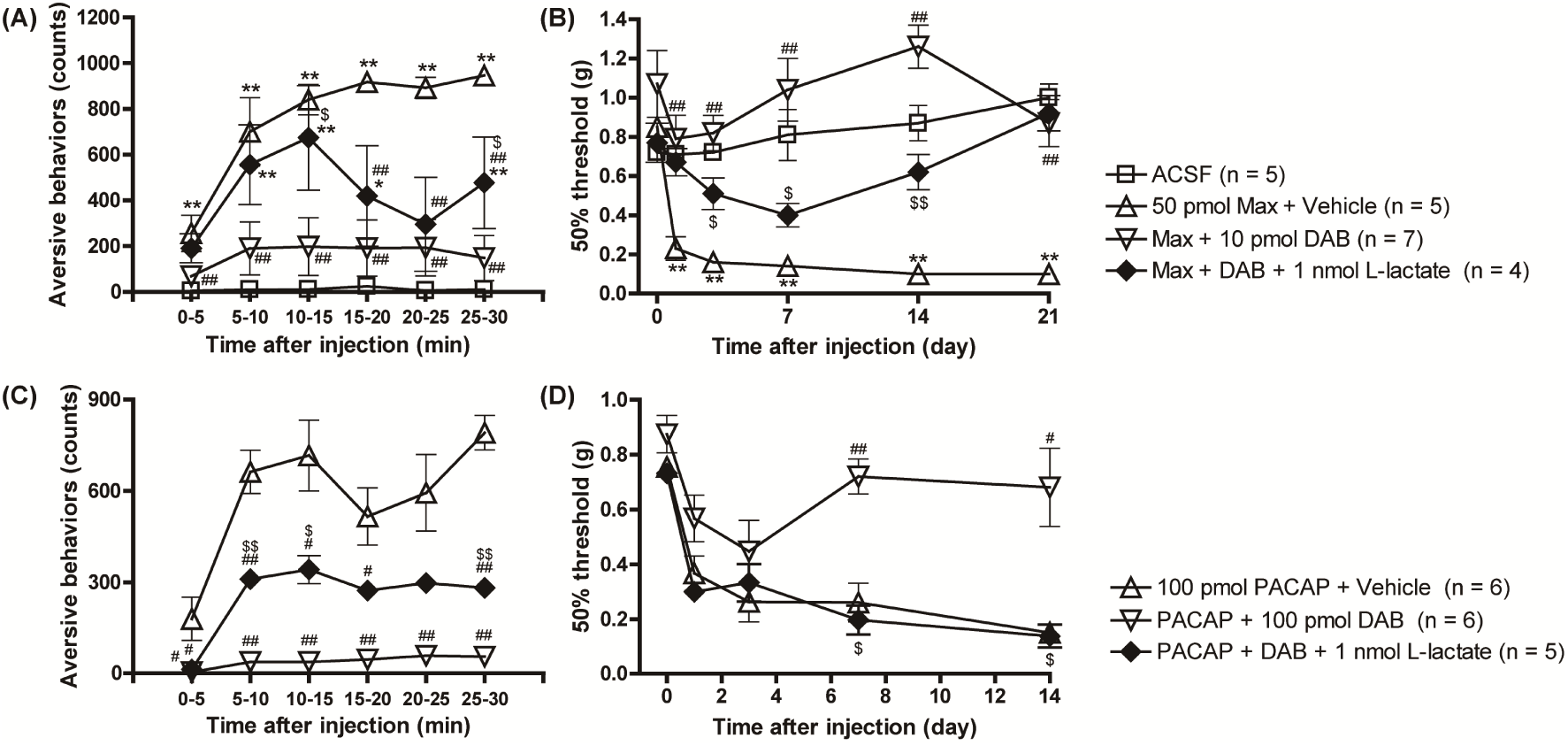
Simultaneous injection of L-lactate reversed the inhibitory effects of DAB on the PAC1 receptor-induced nociceptive behaviors. I.t. co-administration of L-lactate (1 nmol) with DAB (10 or 100 pmol) reinstated Max (**A, B**)- and PACAP (**C, D**)-induced aversive behaviors (**A, C**) and mechanical allodynia (**B, D**). *P<0.05 and **P<0.01 when compared with ACSF data. ^#^P<0.05 and ^##^P<0.01 when compared with Max + Vehicle in (A) and (B) or PACAP + Vehicle in (C) and (D). ^$^P<0.05 and ^$$^P<0.01 when compared with DAB-treated data. Statistical significance was evaluated by the Tukey test for (A) and (C), and Steel test for (B) and (D).

### Possible involvement of PKC in the PACAP/PAC1 receptor-evoked glycogenolysis in cultured spinal cord astrocytes

We prepared cultured astrocytes from newborn mice spinal cords, and confirmed that most cells were suggested to be astrocytes, < 1% cells were oligodendrocytes, and no neurons or microglial cells were detected, when visualized with immunocytochemistry for specific markers of individual cells (***Figure 3 - figure supplement 1***). In order to identify the receptor subtypes involved in the PACAP-evoked glycogenolysis in the spinal astrocytes, we exposed PACAP, VIP or Max and determined glycogen amount in the cells. Exposure of PACAP (0.0001 ~ 10 nM) or Max (0.1 ~ 10 nM) dose-dependently decreased glycogen amount (***Figure 3A, B***), but VIP failed to decrease the glycogen amount even at the concentration of 10 nM (***Figure 3C***), suggesting that PAC1 receptor is primarily involved in the glycogenolysis. Interestingly, this pharmacological characteristic of glycogenolysis was very similar to that of the PACAP-induced nociceptive behaviors in mice reported previously (***Ohnou et al., 2016; Yokai et al., 2016***). This PACAP-induced glycogenolysis was completely abolished by the DAB (1 mM) (***Figure 3D***), indicating that the glycogenolysis by PACAP would be mediated by PYGB. Since PAC1 receptor can couple with both G_s_ and G_q_ proteins, leading to the activation of the protein kinase A (PKA) and protein kinase C (PKC), respectively (***Deutsch and Sun, 1992***), we co-applied GF109203X as a PKC inhibitor, H89 or Rp-8-Br-cAMPS as a PKA inhibitor, or Sq22.536 as an adenylate cyclase (AC) inhibitor with PACAP. We found that GF109203X (5 μM) significantly inhibited the PACAP-evoked glycogenolysis (***Figure 3E***), but neither the PKA inhibitors (***Figure 3F***), nor the AC inhibitor (***Figure 3 - figure supplement 2A***) affected the glycogenolysis. Although these results may contradict previous reports that cAMP/PKA signaling pathway stimulated glycogenolysis in chick brain or cultured mouse cortical astrocytes (***Sorg and Magistretti, 1991; Gibbs, 2016***), forskolin, an AC activator, significantly stimulated glycogenolysis in cultured spinal astrocytes (***Figure 3 - figure supplement 2B***). These results suggested that PAC1 receptor might not be coupled with the G_s_ protein/PKA pathway, and support the significance of the G_q_ protein/PKC pathway for the PACAP/PAC1 receptor-induced glycogenolysis in our cultured spinal cord astrocytes.

**Figure 3.**
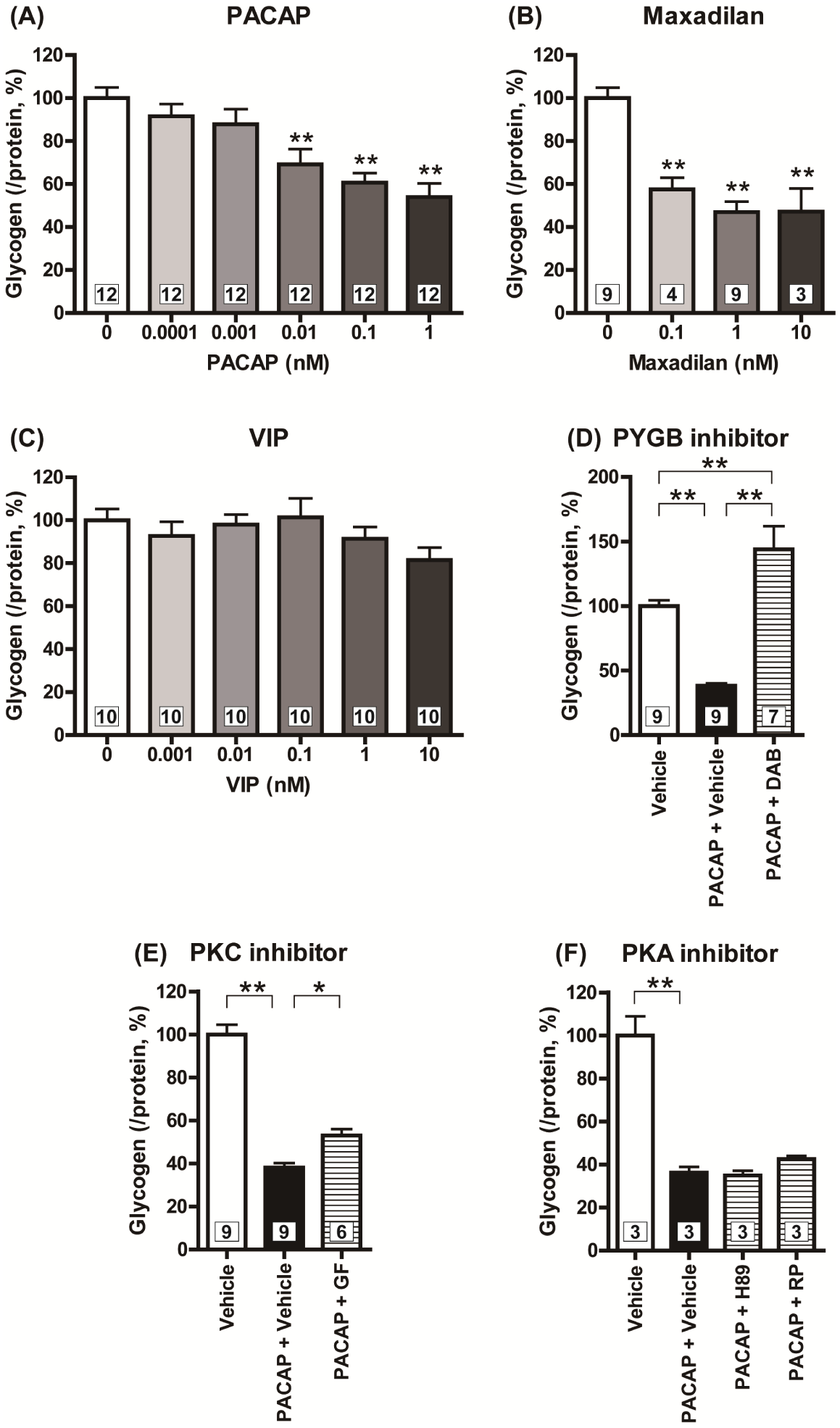
PACAP/PAC1 receptor/PKC signaling pathway induced glycogenolysis in cultured spinal cord astrocytes. Cultured astrocytes were exposed to PACAP (0.0001 ~ 1 nM) (**A**), Max (0.1 ~ 10 nM) (**B**) or VIP (0.001 ~ 10 nM) (**C**), and glycogen amounts contained in the astrocytes were measured 60 min after the exposure. Cultured astrocytes were pre-incubated with DAB (1 mM, **D**), GF109203X (GF, 5 μM, **E**), H89 (10 μM, **F**), or Rp-8Br-cAMPS (RP, 100 μM, **F**) for 30 min prior to PACAP (1 nM) exposure, and glycogen amounts contained in the astrocytes were measured 60 min after the PACAP exposure. *P<0.05 and **P<0.01 when compared with 0 nM (Vehicle) data in (A) ~ (C), and compared groups are indicated above in (D) ~ (F). Statistical significance was evaluated by the Dunnett test for (A) ~ (C), and Tukey test for (D) ~ (F). Exact sample sizes are indicated in the graphs.

### PKC is crucial for the PACAP/PAC1 receptor-evoked lactate secretion in the cultured spinal astrocytes

To investigate whether lactate secretion from astrocytes is increased by the PACAP/PAC1 receptor/PKC pathway, we measured the lactate amounts in the supernatants from the cultured spinal astrocytes after drug treatment. PACAP (1 nM) significantly increased lactate amounts in the supernatant in both 0 ~ 60 min (early phase) and 60 ~ 120 min (late phase) after the exposure (***Figure 4A, B***). PMA (100 nM), a PKC activator, significantly elevated lactate secretion in the late phase (***Figure 4C, D***), while forskolin (5 μM) failed to enhance lactate secretion (***Figure 4E, F***). In addition, GF109203X (5 μM) significantly inhibited the PACAP-activated lactate secretion in both phases (***Figure 4G, H***), suggesting the importance of PKC, but not PKA, in the PACAP-activated lactate secretion. Furthermore, the inhibition of glycogenolysis by DAB significantly decreased the lactate amount in the supernatant from the astrocytes particularly in the late phase (***Figure 4I, J***), indicating that glycogenolysis is also inevitable for the lactate secretion. These results pointed out that PACAP/PAC1 receptor/PKC pathway is a key signaling mechanism for the spinal ANLS activation.

**Figure 4.**
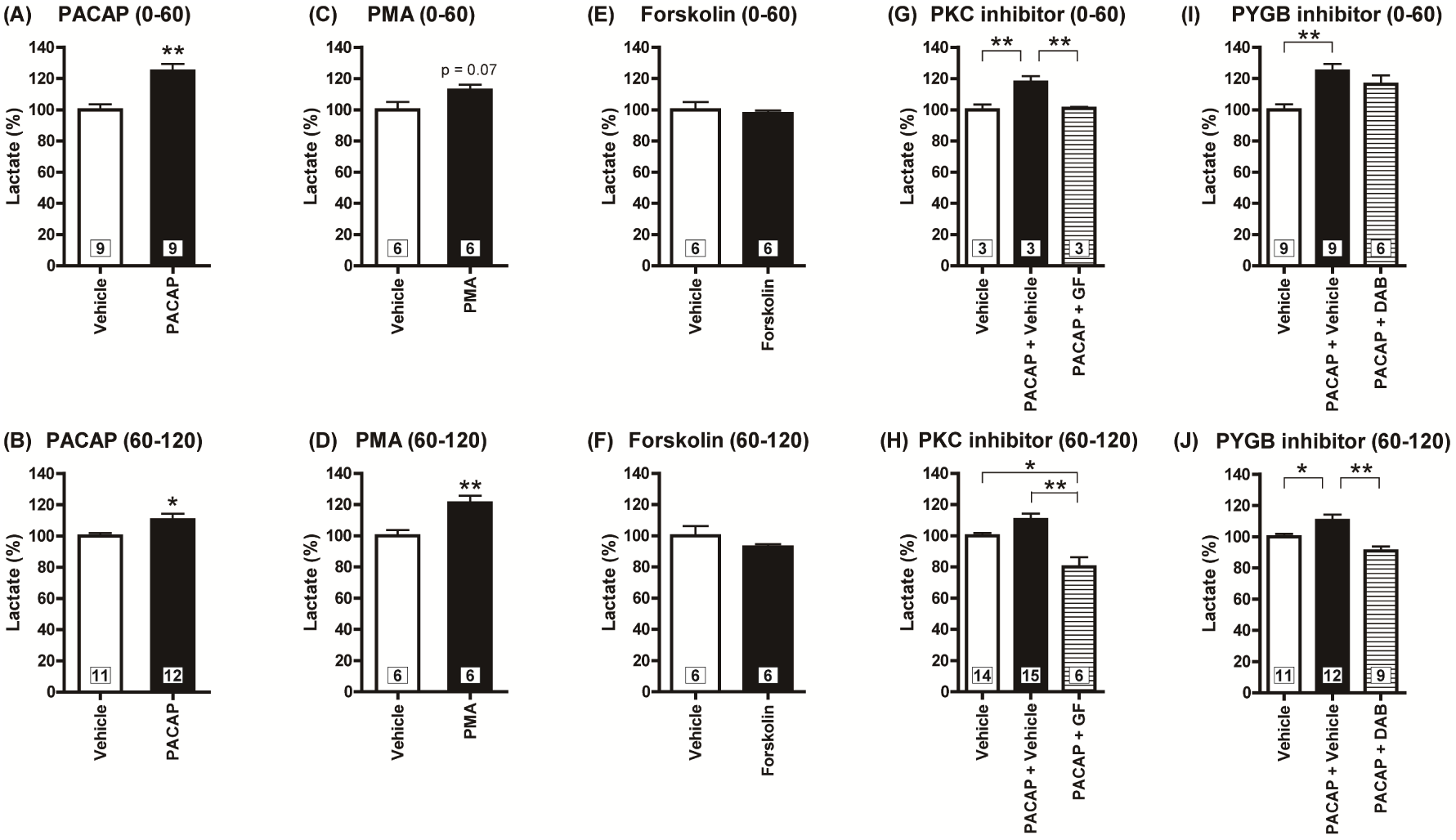
A crucial role of PACAP/PAC1 receptor/PKC-induced glycogenolysis in the lactate secretion from cultured spinal astrocytes. After the treatment with PACAP (1 nM) (**A, B**), PMA (100 nM) (**C, D**) or forskolin (5 μM) (**E, F**), the supernatants from the cultured spinal astrocytes were differentially harvested during 0 ~ 60 min (**A, C, E**) or 60 ~ 120 min (**B, D, F**), and lactate amounts were measured. Cultured astrocytes were pre-incubated with GF109203X (GF, 5 μM, **G, H**) or DAB (1 mM, **I, J**) for 30 min prior to PACAP exposure, and then exposed to PACAP (1 nM) for 120 min. Culture supernatants were differentially harvested during 0 ~ 60 min (**G, I**) or 60 ~ 120 min (**H, J**) after the PACAP addition, and lactate amounts were measured. *P<0.05 and **P<0.01 when compared with vehicle data in (A) ~ (F), and compared groups are indicated above in (G) ~ (J). Statistical significance was evaluated by the Student t-test for (A) ~ (F), and Tukey test for (G) ~ (J). Exact sample sizes are indicated in the graphs.

### Blockade of the PACAP/PAC1 receptor-induced nociceptive behaviors by the PKC inhibitor, GF109203X

In order to investigate whether PACAP/PAC1 receptor-induced nociceptive behaviors were also mediated through PKC pathway, effects of GF109203X was investigated. Both the aversive behaviors and the mechanical allodynia evoked by PACAP were dose-dependently inhibited by the simultaneous injection of GF109203X at a concentration range from 1 to 100 pmol (***Figure 5A, B***), suggesting the importance of PKC activity in the PACAP/PAC1 receptor-induced nociceptive behaviors.

**Figure 5.**
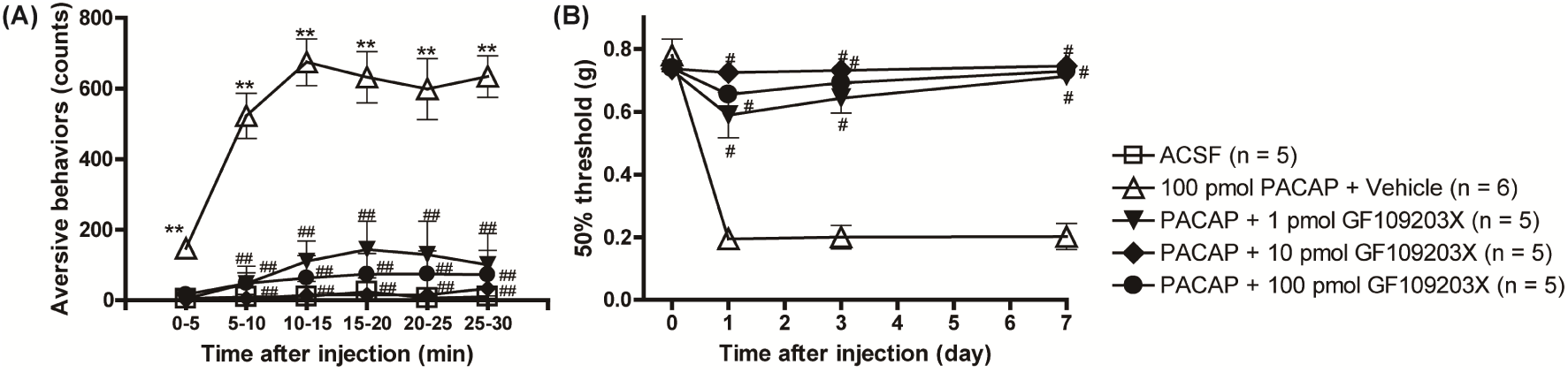
The importance of PKC in the PACAP/PAC1 receptor-induced nociceptive behaviors. Simultaneous i.t. injection of GF109203X (1 ~ 100 pmol) with PACAP (100 pmol) dose-dependently reduced the PACAP-induced aversive behaviors (**A**) and mechanical allodynia (**B**). *P<0.05 and **P<0.01 when compared with ACSF data. ^#^P<0.05 and ^##^P<0.01 when compared with PACAP + Vehicle data. Statistical significance was evaluated by the Tukey test for (A), and Steel test for (B).

### Pharmacological inhibition of monocarboxylate transporters attenuated the PACAP/PAC1 receptor-induced nociceptive behaviors

Finally, we asked whether lactate transport participated in the PACAP/PAC1 receptor-induced nociceptive behaviors. L-lactate is one of the substrates of monocarboxylate transporters (MCTs), and MCTs are found in the brain where three isoforms — MCT1, MCT2 and MCT4 — have been described. Although the distribution patterns of these MCTs are not well-known in the spinal cord, each of these isoforms is suggested to exhibit a distinct regional and cellular distribution in rodent brain. At the cellular level, MCT1 is known to be expressed by endothelial cells of microvessels, by ependymocytes as well as by astrocytes. MCT4 expression appears to be specific for astrocytes. By contrast, the predominant neuronal MCT is suggested to be MCT2 (***Pierre and Pellerin, 2005***). It is reported that AR-C155858 potently inhibits MCT1 and MCT2, but not MCT4 (***Ovens et al., 2010***). In our cultured spinal astrocytes, AR-C155858 (1 μM) significantly inhibited lactate secretion evoked by PACAP at both phases (***Figure 6A, B***). Simultaneous i.t. injection of AR-C155858 (1 nmol) with PACAP significantly inhibited the development of the aversive behaviors (***Figure 6C***). Intriguingly, i.t. administration of AR-C155858 transiently alleviated the mechanical allodynia at 7 days after i.t. injection of PACAP (***Figure 6D***), suggesting that spinal ANLS activation evoked by a single i.t. injection of PACAP may persist at least for 7 days.

**Figure 6.**
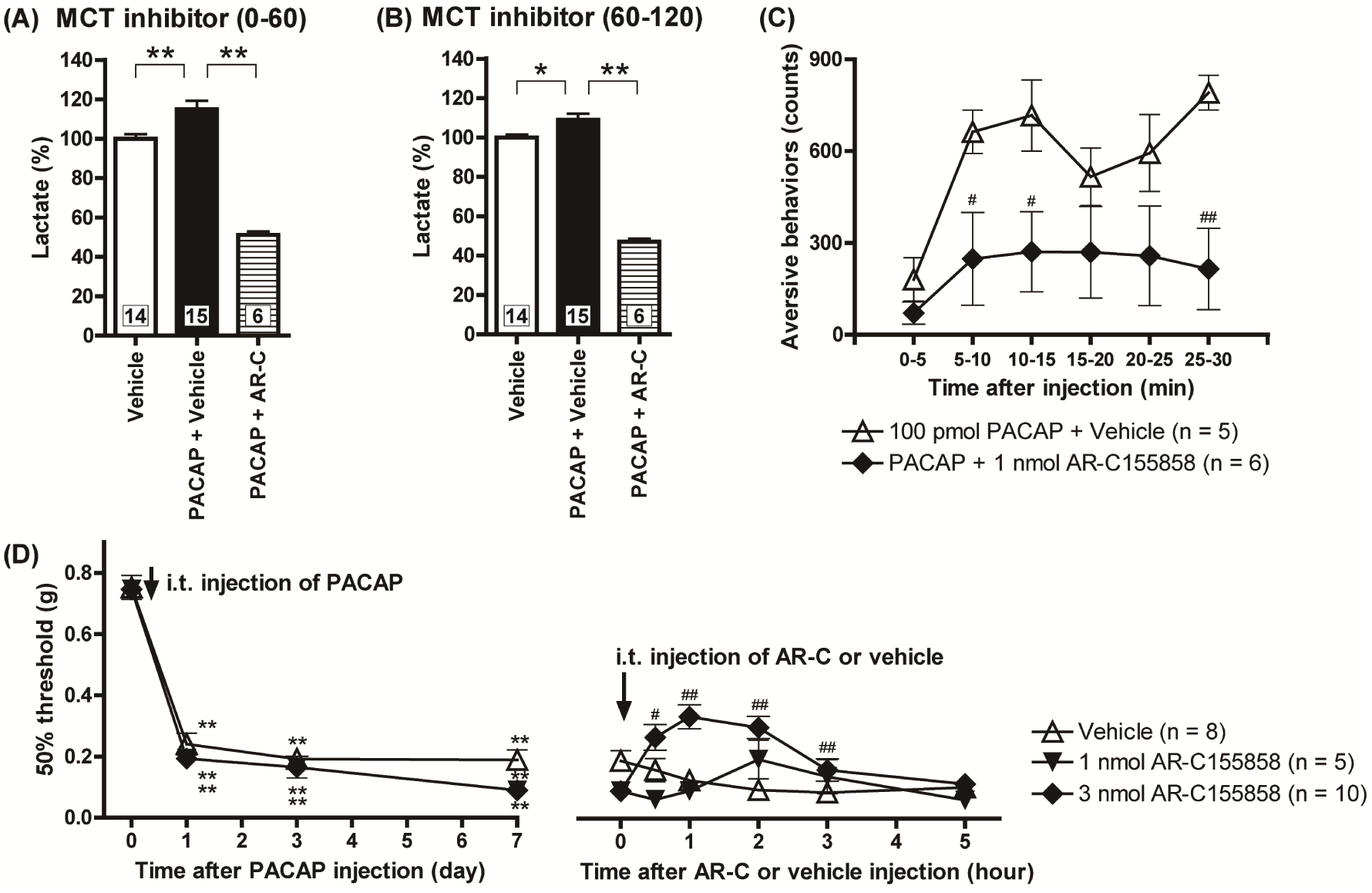
Spinal MCTs play a crucial role in both induction and maintenance of the PACAP/PAC1 receptor-induced nociceptive behaviors. Cultured spinal astrocytes were pre-incubated with AR-C155858 (AR-C, 1 μM) 30 min prior to PACAP exposure, and then exposed to PACAP (1 nM) for 120 min. The culture supernatants were differentially harvested during 0 ~ 60 min (**A**) or 60 ~ 120 min (**B**) after the PACAP addition, and lactate amounts in the culture supernatants were measured. *P<0.05 and **P<0.01 (compared groups are indicated above). Exact sample sizes are indicated in the graphs. (**C**) I.t. co-administration of AR-C155858 (1 nmol) with PACAP reduced the expression of the aversive behaviors. *P<0.05 and **P<0.01 when compared with PACAP + Vehicle data. (**D**) Transient reversal of the PACAP-evoked mechanical allodynia by intrathecal AR-C155858. AR-C155858 was injected 7 days after intrathecal PACAP. **P<0.01 when compared with pre-PACAP data. ^#^P<0.05 and ^##^P<0.01 when compared with vehicle-treated control. Statistical significance was evaluated by the Tukey test for (A) and (B), Student t-test for (C), and Steel test for (D).

## Discussion

In this study, we have further characterized the nociceptive behaviors induced by the spinal PAC1 receptor activation and made the following findings: 1) PACAP could be an endogenous inducer for spinal ANLS activation; 2) spinal ANLS activation by PACAP/PAC1 receptor signaling contributed to nociceptive behaviors such as aversive behaviors and mechanical allodynia; 3) PKC activation (possibly in the spinal astrocytes) played an important role in the PACAP/PAC1 receptor-evoked spinal ANLS; and 4) spinal ANLS activity underling the generation of the nociceptive behaviors persisted at least for 7 days after the PAC1 receptor activation by a single i.t. injection of PACAP.

### PACAP/PAC1 receptor-evoked ANLS activation is crucial for the nociceptive behaviors

In the present study, we showed that the simultaneous i.t. injection of DAB, the inhibitor of glycogenolysis, blocked the development of aversive behaviors and mechanical allodynia evoked by Max or PACAP, and this inhibition by DAB was reversed by the combinational injection of L-lactate. These results suggested that the ANLS activation may have significant role in the nociceptive behaviors by PACAP/PAC1 receptor activation.

Chronic pain may need maladaptive synaptic plasticity in neural circuits (***Sandkuhler, 2007; Kuner and Flor, 2016***). An in vivo electrophysiological study showed that DAB inhibited the long-term potentiation by tetanus stimulus in hippocampal Schaffer collateral-area CA1, and this impairment by DAB was rescued with the exogenous L-lactate (***Suzuki et al., 2011***). Furthermore, L-lactate alone was shown to potentiate AMPA receptor- or NMDA receptor-mediated inward currents, increase intracellular calcium, and also upregulate the markers for neuronal activation (***Suzuki et al., 2011;Nagase et al., 2014; Yang et al., 2014***). Thus, PACAP/PAC1 receptor-evoked spinal ANLS might potentiate AMPA/NMDA receptors-mediated responses and induce long-term potentiation, which might lead to aversive behaviors and long-lasting mechanical allodynia.

Regarding the initiation and maintenance of inflammatory and neuropathic pain, recent progresses also focus on the critical role of astrocytes in the spinal cord and brain (***Ji et al., 2014; Grace et al., 2014***). Previously, we showed that i.t. injection of Max or PACAP induced rapid and prolonged upregulation of spinal GFAP expression level in parallel with the aversive behaviors and long-lasting mechanical allodynia, and simultaneous application of L-α-aminoadipate, an astrocyte toxin, with Max or PACAP markedly suppressed the aversive behaviors and the mechanical allodynia (***Ohnou et al., 2016; Yokai et al., 2016***). Interestingly, it is suggested that glycogen metabolism is associated with maturation of astrocytes, in which the expressional levels of GFAP and glycogen synthase or phosphorylase are increased in parallel (***Brunet et al., 2010***). In accordance with this notion, our present study revealed that ANLS activation indeed contributed to both development and maintenance of PACAP/PAC1 receptor-evoked nociceptive behaviors, since i.t. injection of AR-C155858 could prevent the development of the PACAP-induced aversive behaviors and ameliorate the mechanical allodynia 7 days after PACAP injection. Although, further study is needed to clarify the persistency of ANLS activation and its role in the long-lasting mechanical allodynia, these results suggested the therapeutic potential of the ANLS inhibitors for chronic pain treatments.

### PACAP/PAC1 receptor/PKC signaling pathway played an important role in spinal glycogenolysis

Glycogen contained in cultured spinal cord astrocytes was significantly decreased by the Max or PACAP, but not VIP, exposure. These results suggested that PAC1 receptor, but not VPAC1 nor VPAC2 receptor, was responsible for activating glycogenolysis. In rat atrial myocytes, a PKC inhibitor peptide reduced the PACAP-induced K_ATP_ currents without affecting the VIP-induced K_ATP_ currents (***Baron et al., 2001***), indicating selective coupling of PKC with PAC1 receptor. Indeed, in our present study, PACAP-evoked glycogenolysis was significantly inhibited by the PKC inhibitor, GF109203X, but not by the PKA inhibitors. These results suggested that PAC1 receptor/PKC signaling pathway is important for the glycogenolysis activation.

Previous reports which observed glycogenolysis in chick brain or cultured mouse cortical astrocytes suggested that cAMP/PKA signaling pathway phosphorylated glycogen phosphorylase and evoked glycogenolysis (***Sorg and Magistretti, 1991; Gibbs, 2016***). But in our cultured spinal astrocytes, both PKA inhibitors and AC inhibitor failed to prevent the PACAP-evoked glycogenolysis. Although the reason why PACAP-evoked glycogenolysis is not sensitive to the PKA inhibitors is currently unknown, this may be due to the absence of the coupling of PACAP/PAC1 receptor to Gs protein in the spinal astrocytes, because both PKA and adenylate cyclase inhibitor failed to inhibit the PACAP-evoked glycogenolysis, and forskolin, a direct AC activator, successfully evoked glycogenolysis. But further study is required to verify this possibility.

It is previously suggested that VIP neurons exert their glycogenolytic action in the neocortical region (***Magistretti and Allaman, 2015***), but in our cultured spinal astrocytes, VIP failed to evoke glycogenolysis at the concentration up to 10 nM. This discrepancy might be due to the regional difference, because in our unpublished results with using cultured cortical astrocytes, VIP (EC_50_ = 0.43 nM) as well as PACAP (EC_50_ = 0.0084 nM) could evoke glycogenolysis. In addition, cortical astrocytes express VPAC2 receptors more abundantly than PAC1 receptor (our unpublished observation). These results support the notion of the regional difference of astrocytic function, and the glycogenolysis of the spinal astrocytes might be primarily mediated by PACAP signaling, while that in the cortical astrocytes might be mediated by both PACAP and VIP signaling.

The reason why DAB treatment with PACAP increased glycogen level more than vehicle control might be due to the constitutive activity of PYGB. However, further research is also necessary to elucidate the basal activity of PYGB in the spinal astrocytes.

### PACAP/PAC1 receptor/PKC signaling-evoked glycogenolysis played a critical role in the lactate secretion

PACAP and PMA significantly enhanced lactate secretion. In addition, GF109203X and DAB significantly inhibited the PACAP-enhanced lactate secretion. These results proposed that PKC and a downstream glycogenolysis were important in the PACAP-enhanced lactate secretion. It is previously reported that the PKC activation resulted in an increase in the levels of MCT1 and MCT4 and the release of lactate in an in vitro skeletal muscle cells (RD cells), while the PKA activation resulted in a significant decrease in MCT1 expression in the RD cells as well as endothelial cells (***Narumi et al., 2010;Narumi et al., 2012; Smith et al., 2012***). Although the PKA activation by dbcAMP (cAMP analogue) or isoproterenol (selective β agonist) was shown to significantly decrease glycogen level in astrocytes/neurons mixed culture, the secretion of lactate, contrary to expectation, was unchanged or decreased (***Tarczyluc et al., 2013***). Consistent with these reports, our preliminary observation using cultured cortical astrocytes, forskolin significantly attenuated the lactate secretion, even though significant glycogenolysis was observed (unpublished observation). These results indicate that increased lactate secretion by PKC might involve the activation of MCT1 and MCT4, and in some cases, PKA might counteract PKC-induced action on lactate secretion. Since forskolin treatment did not stimulate lactate secretion in the spinal astrocytes, PACAP/PAC1 receptor/PKC signaling, but not PKA signaling, would be involved in the lactate secretion in our present experimental condition.

It may be noteworthy that DAB significantly inhibited the lactate secretion in the late phase after PACAP exposure, but failed to inhibit in the early phase. Our results may be consistent with a previous report that DAB significantly attenuated the lactate secretion during 60 ~ 180 min after the hypoglycemia on the astrocytes/neurons mixed culture, while equivalent amount of lactate was secreted irrespective of the presence of DAB during 0 ~ 60 min, even DAB completely inhibited glycogenolysis at both timings (***Tarczyluk et al., 2013***). Thus, these observations may indicate that the secreted lactate in the late phase is originated from glycogenolysis, and the secretion in the early phase may be partly derived from the readily releasable pool of lactate in the spinal astrocytes.

### PACAP/PAC1 receptor-induced PKC activation is important for the nociceptive behaviors

We found that developments of the PACAP/PAC1 receptor-induced aversive behaviors and mechanical allodynia were significantly attenuated by the PKC inhibition with GF109203X. Although many lines of evidence strongly suggested the significance of PKC in inflammatory or neuropathic pain (***Velázquez et al., 2007***), it is still unknown what kinds of PKC isozymes actually contribute to the PACAP/PAC1 receptor-evoked ANLS activation, or in which cell types the PKC was activated. PKC has several isozymes and, western blotting showed that phospho-PKC-δ, -θ and -ζ levels were significantly increased in ipsilateral spinal cord after spinal nerve injury, but expression levels of phospho-PKC-α and -β were not significantly changed (***Gosselin et al., 2013***). Further double labeling immunofluorescence showed that phospho-PKC-δ co-localized with astrocytic markers and phospho-PKC-θ and -ζ co-localized with neuronal markers in spinal cord (***Gosselin et al., 2013***). Another research group reported that the i.t. injection of a PKC-δ inhibitor significantly attenuated paclitaxel-induced mechanical allodynia in mice (***He and Wang, 2015***). These results suggested that PKC-δ might be an astrocytic PKC isoform and contribute to the pain behaviors observed in this study. Interestingly, it is indicated that PMA-induced stimulation of MCT4 expression can be mediated through a novel PKC isozyme, especially PKC-δ, and rottlerin, a selective PKC-δ inhibitor, cancelled PMA-induced MCT4 protein expression in the RD cells (***Narumi et al., 2012***). Further study is needed to elucidate the contribution of PKC isozymes in the PACAP/PAC1 receptor-induced spinal ANLS and nociceptive behaviors.

It may be worth noting our previous data here that i.t. injection of the PKA inhibitor, Rp-8-Br-cAMPS, could ameliorate the PACAP-induced aversive behaviors (***Ohnou et al., 2016***). Currently it may be difficult to precisely reconcile with the present observation that the PKA inhibitor failed to prevent PACAP-evoked glycogenolysis in the spinal astrocytes, but we could speculate that neuronal PKA signaling would also contribute to the expression of the pain behaviors. However, further extensive studies are necessary to uncover the interaction of PKC and PKA signaling in the PACAP/PAC1 receptor-induced pain behaviors.

### MCTs had an important role in PACAP/PAC1 receptor-induced nociceptive behaviors

In cultured spinal astrocytes, AR-C155858 potently inhibited PACAP-evoked lactate secretion. In the behavioral study, the simultaneous i.t. injection of AR-C155858 significantly attenuated the development of the PACAP-induced aversive behaviors. In addition, AR-C155858 also transiently reversed the PACAP-induced mechanical allodynia 7 days after the PACAP injection. These observations suggested the important contribution of lactate transportation via MCTs to the induction as well as the maintenance of the nociceptive behaviors evoked by the spinal PACAP/PAC1 receptor signaling. The significance of MCTs on the synaptic plasticity was previously reported. Hippocampal MCT1 expression but not MCT2 or MCT4 expression was significantly upregulated upon inhibitory avoidance training in rats, and blocking the function of MCT1, MCT2 or MCT4 by the antisense oligonucleotides impaired long-term memory (***Suzuki et al., 2011***). Moreover, in the cocaine-induced conditioned place preference test, re-exposure to a cocaine-associated context significantly increased the expression of MCT1 and MCT2, and disrupting their expression led to the impairment of reconsolidation (***Zhang et al., 2015***). At the cellular level, L-lactate-induced increased expressions of immediately early genes such as *Arc*, *c-fos* and *Zif268* were blocked in the presence of a MCT inhibitor, UK5099 (***Yang et al., 2014***). To our knowledge, the present study is the first report establishing the significance of MCTs in nociceptive behaviors.

In summary, we have shown that the interaction between spinal dorsal horn neurons and astrocytes evoked by PACAP/PAC1 receptor-induced ANLS activation is critically involved in the development and maintenance of the nociceptive behaviors. The signaling pathway linking between PAC1 receptor activation and ANLS might be at least partially mediated by PKC pathway. Although verifying the possible involvement of PACAP/PAC1 receptor signaling in human pain as well as animal pain models requires further rigorous studies, targeting spinal ANLS may provide a new opportunity to treat intractable chronic pain.

## Materials and methods

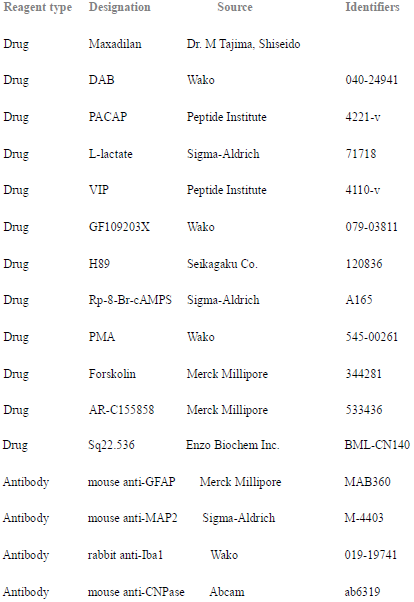
Key resources table.

### Animals

Male ddY mice (6 ~ 12 weeks old) or pregnant female ddY mice were purchased from Kyudo Co. Ltd. (Kumamoto, Japan) and housed under controlled temperature (24 ± 1°C) and humidity (55 ± 10%) with a 12-h light/dark cycle with food and water freely available.

### Ethics

The animal experiments were approved by the Animal Care Committee of Kagoshima University (approval no. MD15082), and were conducted in accordance with the ethical guidelines for the study of experimental pain in conscious animals of the International Association for the Study of Pain.

### Intrathecal (i.t.) injection and behavioral observation

I.t. injection was given in a volume of 5 μl by percutaneous puncture through an intervertebral space at the level of the fifth or sixth lumbar vertebra, according to a previously reported procedure (***Yokai et al., 2016***). An investigator, who was unaware of the drug treatment, performed all of the behavioral experiments.

Aversive behavior was evaluated according to our previous report (***Ohnou et al., 2016***). Before i.t. injection, mice were placed and habituated in a glass cylinder (φ14 × 18 cm) with a filter paper at the bottom over 20 min. Immediately after i.t. injection, the mice were placed again in the same glass cylinder and the number of pain behaviors consisting of licking, biting and scratching directed toward the caudal part of the body was counted every 1 min. The cumulative number of events was pooled over 5 min bins of observation, and analyzed.

The assessment of mechanical thresholds was also carried out according to the previously described methods (***Yokai et al., 2016***). Briefly, mechanical sensitivity was evaluated with calibrated von Frey hairs (Stoelting, Wood Dale, IL) by measuring the tactile stimulus producing a 50% likelihood of hind paw withdrawal response (50% gram threshold), which was determined using the up-down paradigm (***Chaplan et al., 1994***).

### Drugs

PACAP (38 amino acid form) and VIP were purchased from Peptide Institute Inc. (Osaka, Japan). Maxadilan was kindly donated by Dr. M Tajima (Shiseido, Japan). 1,4-dideoxy-1,4-imino-d-arabinitol (DAB), GF109203X and phorbol 12-myristate 13-acetate (PMA) were obtained from Wako Pure Chemical Industries, Ltd. (Osaka, Japan). Forskolin and AR-C155858 was from Merck Millipore (Darmstadt, Germany). H89 was from Seikagaku Co. (Tokyo, Japan). L-lactate and Rp-8-Br-cAMPS were from Sigma-Aldrich Co. LLC (St. Louis, MO). Sq22.536 was from Enzo Biochem Inc. (Farmingdale, NY). These drugs were made up as concentrated stock solution in MilliQ water, 0.1 M acetic acid or dimethyl sulfoxide, aliquoted, and stored at –30°C and diluted just before use. Aliquot of drugs were diluted to the desired concentration in artificial cerebrospinal fluid (ACSF: NaCl 138 mM, KCl 3 mM, CaCl_2_ 1.25 mM, MgCl_2_ 1 mM, D-glucose 1 mM) immediately prior to use. The doses for i.t. injection of PACAP38 (100 pmol) and maxadilan (50 pmol) we chose in this study were determined according to our previous reports (***Shimizu et al., 2004;Ohnou et al., 2016; Yokai et al., 2016***).

### Cultured spinal cord astrocytes

Spinal cords from postnatal day 1 and 2 mice were dispersed by 0.25% trypsin and subsequently plated on culture flasks which were previously coated with 0.75 ug/mL poly-l-lysine (Sigma-Aldrich, St. Louis, MO). Cells were cultured in Dulbecco’s modified Eagle medium (Nacalai Tesque, Osaka, Japan) supplemented with 10% fetal bovine serum (Hyclone, South Logan, UT) until cell density reached to confluent. After reaching to confluent, culture flasks were shaken at 240 rpm for 6 hrs to remove microglia or oligodendrocyte precursor cells. Remaining astrocytes on culture flasks were then detached and subsequently plated on the appropriate culture dishes at the density of 1 × 10^4^ cells/cm^2^, and used when the cell number reached to the confluent again, which usually took 3 days. All of the cultured astrocytes used in the current experiments was passaged by 3 or 4 times.

### Glycogen assay

Cultured spinal astrocytes were exposed to PACAP, VIP, Max or forskolin for 1 hr, washed in the ice-cold phosphate buffered saline 3 times, and harvested in ice-cold MilliQ water by scratching. Samples were then boiled for 10 min, sonicated, and centrifuged. Glycogen amount in the supernatant was measured by the Glycogen Assay Kit (BioVision Inc., San Francisco, CA) according to the manufacturer’s instruction. The exposure of DAB, H89, Rp-8-Br-cAMPS, Sq22.536 or GF109203X was started 30 min before PACAP exposure. The amount of glycogen was normalized by the amount of the protein contained in the samples, which was measured by the Bradford’s assay kit (Bio-Rad, Hercules, CA) according to the manufacturer’s instruction.

### Lactate assay

Culture medium was replaced to Krebs-Ringer buffer (NaCl 135 mM, KCl 5 mM, CaCl_2_ 1 mM, MgSO_4_ 1 mM, KH_2_PO_4_ 0.4 mM, D-glucose 5.5 mM, HEPES 20 mM), and the astrocytes were habituated in this buffer for 2 hrs. The cultured spinal astrocytes were then exposed to PACAP, PMA or forskolin for 60 min, and conditioned buffer was harvested (0 ~ 60 min). Subsequently, flesh Krebs-Ringer buffer with drugs was added on the cultured astrocytes again, and harvested the conditioned buffer for another 60 min (60 ~ 120 min). The time course was determined according to the previous paper which suggested DAB inhibited lactate secretion in 60 ~ 180 min after the treatment, but not in 0 ~ 60 min (***Tarczyluk et al., 2013***). Lactate amount in the conditioned Krebs-Ringer buffer was measured by the L-lactate Assay Kit (BioVision Inc.) according to the manufacturer’s instruction. The exposure of DAB or GF109203X was started 30 min before PACAP treatment.

### Statistical analysis

Experimental data are expressed as mean ± SEM. For mechanical threshold analyses, we employed the Mann-Whitney U-test for single comparisons or the Friedman test followed by the Steel test for multiple comparisons. For other analyses, single comparisons were made using the Student two-tailed unpaired *t*-test, and for multiple comparisons, one-way analysis of variance followed by the Dunnett or Tukey test was used. P<0.05 was considered statistically significant.

## Acknowledgements

The authors thank all the staff members of the Institute of Laboratory Animal Science Reserch Support Center, Kagoshima University.

## Additional information

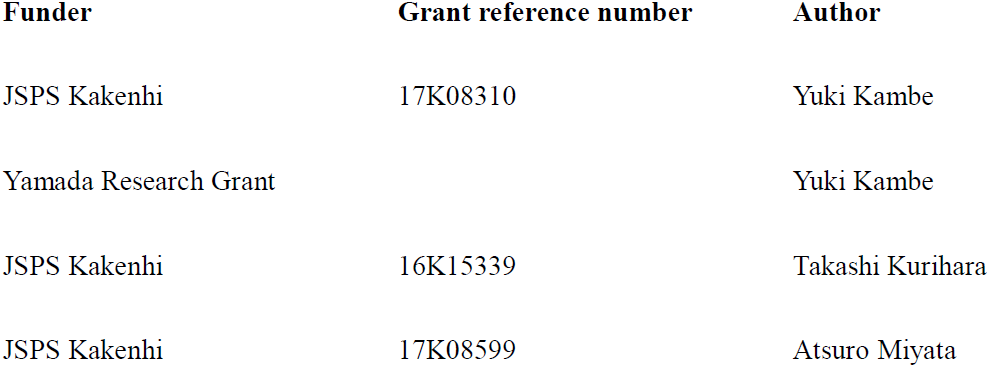
Funding.

### Author contributions

YK carried out experiments, performed statistical analysis, and drafted the manuscript. MY carried out experiments and performed statistical analysis. TK conceived, participated in the design of the study, performed behavioral studies, and wrote the manuscript. AM and IT participated in the design of the study and reviewed the manuscript. All authors read and approved the final manuscript.

---

**Figure 3 - figure supplement 1.**
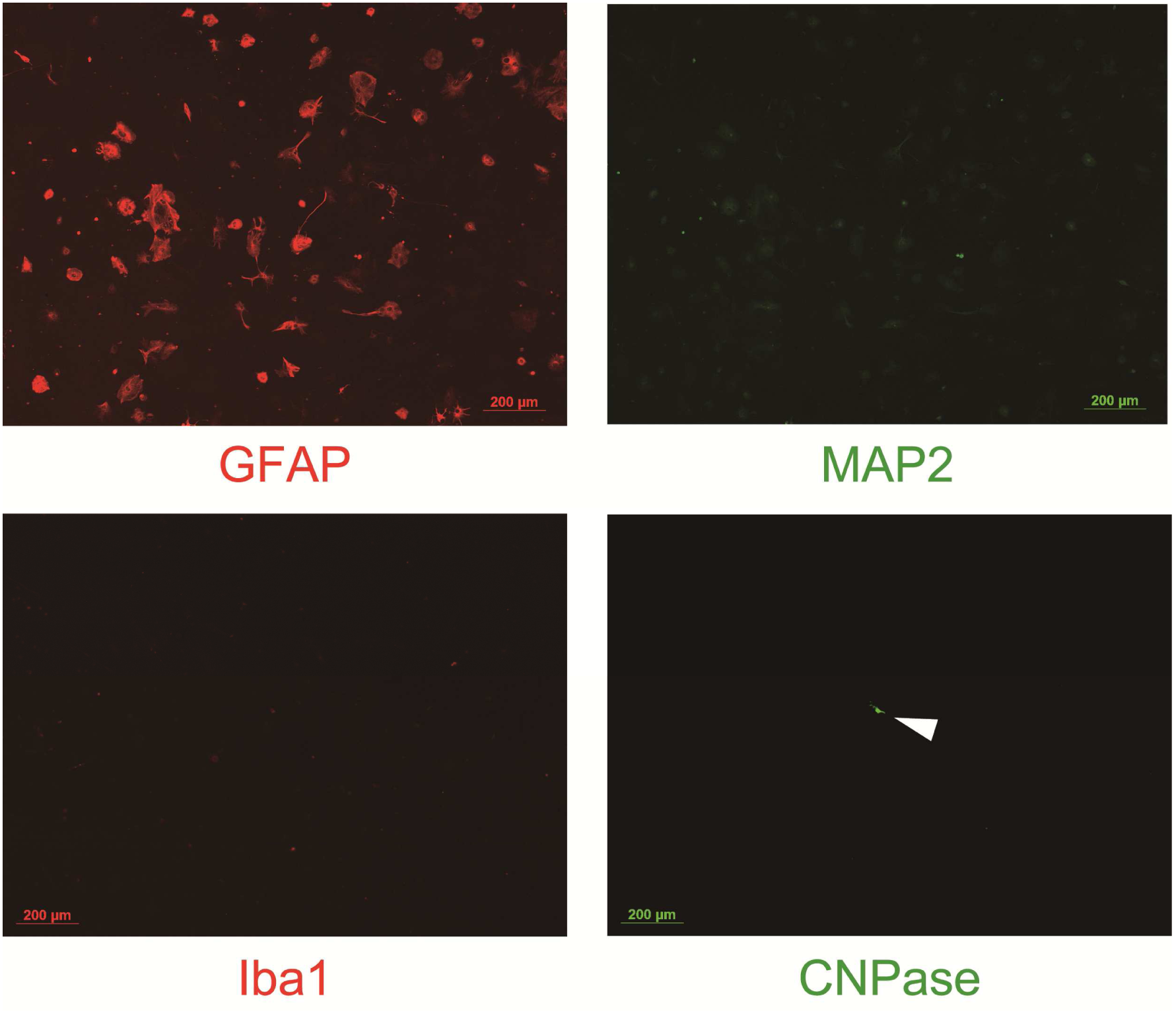
Representative immunofluorescent images of the cultured spinal astrocytes. The purity of astrocytes was examined by staining with primary antibodies against GFAP (an astrocyte marker), MAP2 (a neuron marker), Iba 1 (a microglia marker) and MBP (an oligodendrocyte marker). Each marker was visualized by an appropriate fluorescent dye-conjugated secondary antibody. Arrowhead represented an incidentally contaminated oligodendrocyte. Bar = 200 μm.

**Figure 3 – figure supplement 2.**
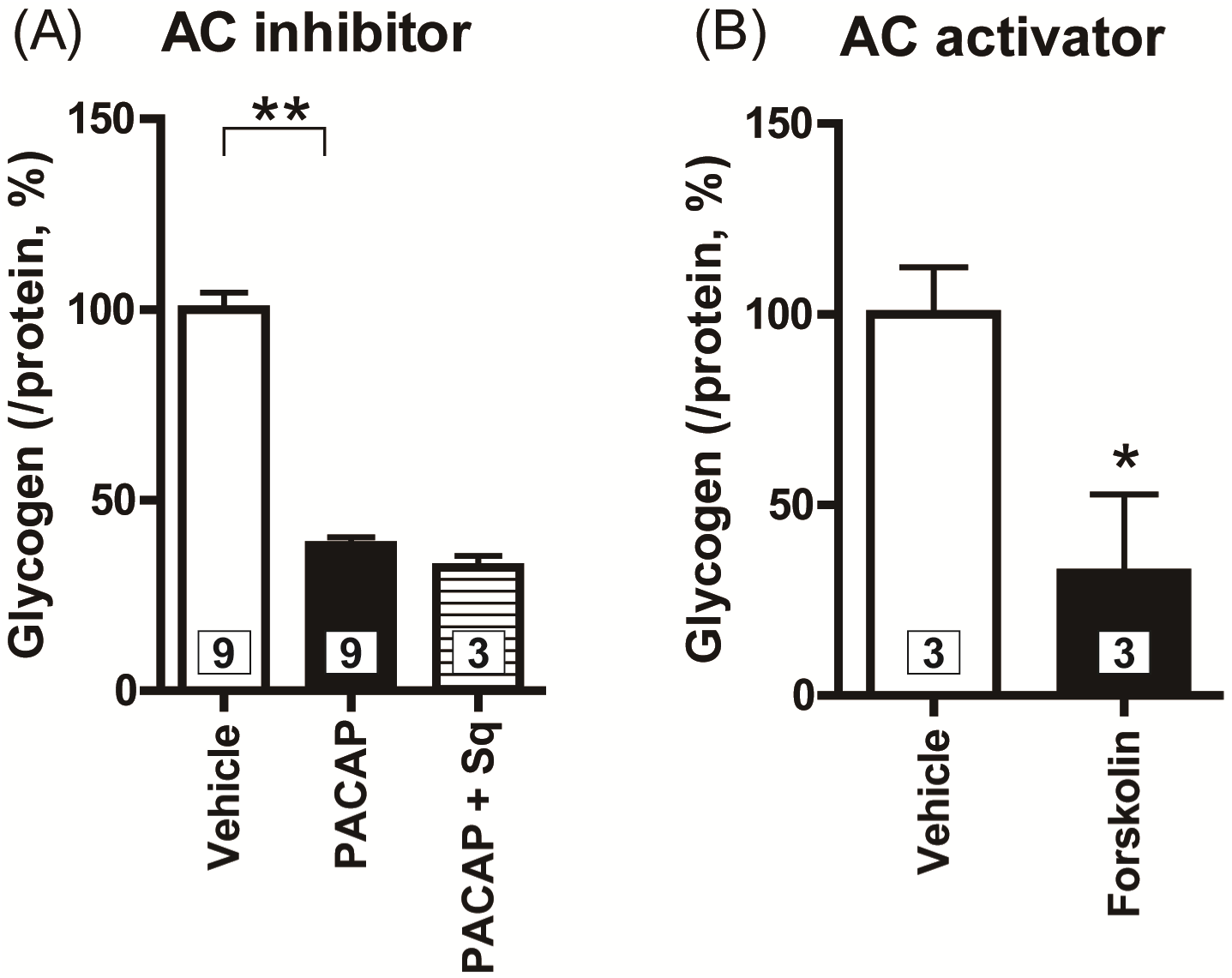
PAC1 receptor might not be coupled with the G_s_ protein/PKA pathway in the cultured spinal astrocytes. Cultured spinal astrocytes were pre-incubated with Sq22.536 (Sq, 100 μM) for 30 min prior to PACAP (1 nM) exposure, and glycogen amounts contained in the astrocytes were measured 60 min after the PACAP exposure (**A**). Cultured spinal astrocytes were exposed to forskolin (5 μM), and glycogen amounts contained in the astrocytes were measured 60 min after the exposure (**B**). *P<0.05 and **P<0.01 when compared with Vehicle. Statistical significance was evaluated by the Dunnett test for (A) and Student t-test for (B). Exact sample sizes are indicated in the graphs.

## Abbreviations used

(ACSF): artificial cerebrospinal fluid
(AC): adenylate cyclase
(ANLS): astrocyte-neuron lactate shuttle
(CNS): central nervous system
(GFAP): glial fibrillary acidic protein
(i.t.): intrathecal
(Max): maxadilan
(MCT): monocarboxylate transporter
(PACAP): pituitary adenylate cyclase-activating polypeptide
(PKA): protein kinase A
(PKC): protein kinase C
(PMA): phorbol 12-myristate 13-acetate
(PYGB): glycogen phosphorylase
(VIP): vasoactive intestinal polypeptide

